# The metallophore staphylopine is essential for survival of *Staphylococcus epidermidis* in human synovial fluid

**DOI:** 10.1101/2025.08.18.670852

**Authors:** Leanne Sims, Muhammad Yasir, Emma Manners, Claire Hill, Heather Felgate, John Wain, Iain McNamara, Mark A Webber

## Abstract

Due to extended life expectancies, prosthetic joint infections are an increasing burden on healthcare institutions worldwide. The most commonly isolated causative agents are staphylococci, though the mechanisms underpinning survival and proliferation in synovial fluid are still not fully understood. In this study, we aimed to identify genes important for survival in synovial fluid using a transposon mutant library and RNAseq.

We produced a transposon mutant library, containing approximately 57,000 unique insertion mutants, in *Staphylococcus epidermidis* strain 846. This library was grown in Müller Hinton broth or processed human synovial fluid samples, and Transposon-directed Insertion Sequencing (TraDIS) was used to identify genes involved in survival in synovial fluid. This identified importance of the his, pur and cnt operons. These genes were also upregulated in both *Staphylococcus epidermidis* 846 and the model *Staphylococcus epidermidis* strain RP62A when exposed to human synovial fluid. All these key pathways contribute to production of the metallophore staphylopine.

To confirm staphyopine production is essential for survival in synovial fluid, a defined transposon insertion in the gene encoding for staphylopine export (*cntE*) mutant was used. This demonstrated impaired survival in synovial fluid compared to the wild type. RT-qPCR also showed that cntE was more highly expressed after exposure to infected synovial fluid (where metals will be depleted) than non-infected fluid.

In conclusion, TraDIS and RNASeq both identified the importance of staphylopine for survival in human synovial fluid. This suggests an opportunity for exploitation for therapeutic or diagnostic use.

**Author Summary:** *Staphylococcus epidermidis* is a common cause of prosthetic joint infection, however accurate diagnosis remains difficult. In this work we explored the genetic basis of *Staphylococcus epidermidis* survival in human synovial fluid using a large transposon mutant library, and identified which genes were differentially expressed upon exposure to the fluid. We found crossover between the datasets, which pointed to the importance of the metal acquisition compound staphylopine. The expression levels of the gene required for staphylopine export were shown to be dependent on the infection status of the individual samples were obtained from. We also found the gene cluster to be conserved in a range of staphylococcal species isolated from cases of prosthetic joint infection. Our work provides a valuable resource in the from of a large transposon mutant library, and provides a greater understanding of the requirements for staphylococcal survival in human synovial fluid, providing potential biomarkers for future diagnostic development.

## Introduction

As populations age total joint arthroplasty becomes a growing burden on healthcare institutions worldwide (1). The increasing incidence of joint replacements is reflected by a commensurate rise in rates of reported prosthetic joint infection (PJI), a complication following 1-2% of primary joint replacement surgeries (2). Treatments for PJI are varied, ranging from debridement and implant retention (DAIR) through single stage revision surgery to the current gold-standard, two-stage revision operations. These treatments are both invasive and costly, and represent an important economic burden; the average hip replacement due to infection was reported in 2019 as costing £50,000 (3).

The majority of PJI cases are caused by *Staphylococcus* species, with *S. aureus* accounting for 22-33% of cases, and coagulase-negative staphylococci (CoNS) 19-41% (4, 5). Despite being the most common cause of PJI, we know little about how staphylococci grow in synovial fluid. When exposed to human synovial fluid, *Staphylococcus aureus* and *epidermidis* rapidly form aggregates (6, 7) which can attached to prostheses possibly leading to an environment protected from antibiotics (8, 9). The mechanisms underpinning aggregate formation and the survival of CoNS in human synovial fluid have not yet been elucidated. Understanding how pathogens cause PJI is needed to develop better ways to prevent or treat PJI.

Diagnosis of infection by CoNS species is complicated, as they are ubiquitous members of the skin microbiome, and differentiation of contaminating commensal versus pathogen often remains impossible (8-10). Longer time to diagnosis impacts on the treatments available, and established biofilm infections by CoNS are more difficult to treat (8, 9), resulting in treatment which is more invasive, requires longer recovery times and incurs much higher costs (11).

Previously, we used machine learning to identify genes important for biofilm formation in a panel of 385 CoNS (12). This work identified that separate groups of *Staphylococcus* have evolved to form biofilms in different ways. One major distinction was between whether strains used a polysaccharide based matrix (encoded by the *ica* operon), or a proteinaceous matrix. We selected one *ica*-positive and one *ica*-negative strain of *S. epidermidis* for this study; RP62A and 846 respectively, to reflect this diversity. To identify genes of importance within these strains, we have combined the use of RNASeq and Transposon Directed Insertion Sequencing (TraDIS). RNASeq was used to identify which genes were expressed from the two strains after exposure to human synovial fluid. In parallel, a TraDIS library produced in *S. epidermidis* 846 was grown in synovial fluid and compared to growth in media to identify key genes required for survival in human synovial fluid. Both these techniques together hold the potential to identify novel targets that can serve in new diagnostic tests or treatments.

## Results

### Production of a transposon mutant library in *Staphylococcus epidermidis* 846, and identification of genes important for growth in human synovial fluid

A transposon mutant library was produced in the *ica*-negative *Staphylococcus epidermidis* 846, using a Tn5 transposon carrying an erythromycin resistance cassette. This library was found to contain approximately 57,000 unique insertion mutants, equating to an average insertion every 50 bp (corresponding to approximately 20 independent mutants per gene). A total of 314 genes were found to be involved in the survival of *Staphylococcus epidermidis* 846 in synovial fluid compared to growth in Mueller-Hinton broth (q<0.05, LFC >1) (Supplementary File S1). Of these, 117 genes were protected during growth in human synovial fluid (reduced insertions due to mutants failing to compete), and 197 genes were found to have more insertions within the gene during growth in human synovial fluid, suggesting disruptions gave a benefit to survival.

The roles of these important genes were studied, and connecting metabolic pathways identified. Insertions across the *pur* operon (*purDHNMFLQSCKE*) (Figure 1) were found to confer a survival benefit to *S. epidermidis* 846. These genes allow the de novo biosynthesis of purines. Mutations in genes which encode proteins within the TCA cycle (*fumC, sucD, sucC* and *acnA*) and central respiration (*pdhD, sucA* and *sucB*) also appeared to confer a survival benefit (Table 1).

**Table 1:** Key pathways which were enriched in the genes identified as playing a role in survival in synovial fluid from the TraDIS dataset.

| KEGG pathway | Pathway description | Observed gene count | Genes in pathway | Strength | Signal | False discovery rate |
| --- | --- | --- | --- | --- | --- | --- |
| ser01110 | Biosynthesis of secondary metabolites | 67 | 246 | 0.33 | 0.7 | 2.47E-06 |
| ser01100 | Metabolic pathways | 104 | 480 | 0.23 | 0.61 | 3.17E-06 |
| ser01200 | Carbon metabolism | 27 | 80 | 0.42 | 0.55 | 0.00094 |
| ser00230 | Purine metabolism | 19 | 47 | 0.5 | 0.55 | 0.0017 |
| ser00020 | Citrate cycle (TCA cycle) | 13 | 27 | 0.58 | 0.5 | 0.0051 |
| ser01120 | Microbial metabolism in diverse environments | 35 | 138 | 0.3 | 0.41 | 0.0051 |
| ser00260 | Glycine, serine and threonine metabolism | 13 | 29 | 0.55 | 0.47 | 0.0063 |
| ser01230 | Biosynthesis of amino acids | 27 | 102 | 0.32 | 0.38 | 0.0101 |
| ser00630 | Glyoxylate and dicarboxylate metabolism | 10 | 21 | 0.57 | 0.41 | 0.0157 |
| ser00640 | Propanoate metabolism | 10 | 24 | 0.52 | 0.34 | 0.0307 |
| ser01210 | 2-Oxocarboxylic acid metabolism | 9 | 20 | 0.55 | 0.35 | 0.0307 |
| ser00620 | Pyruvate metabolism | 14 | 44 | 0.4 | 0.32 | 0.0322 |
| ser00280 | Valine, leucine and isoleucine degradation | 7 | 13 | 0.63 | 0.35 | 0.0341 |

**Figure 1:**
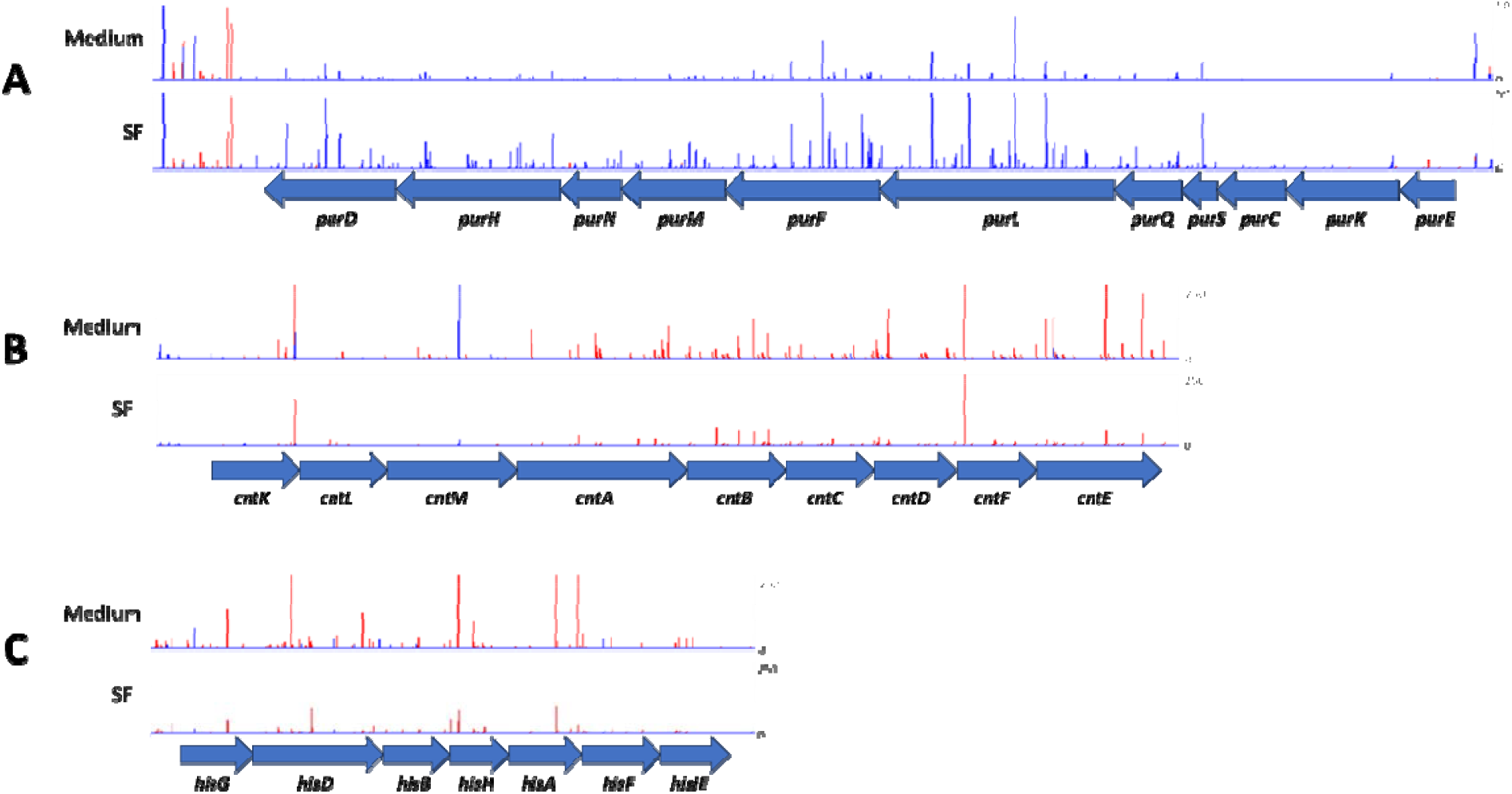
Differential abundance of *S. epidermidis* transposon insertion mutants after growth in synovial fluid. Peaks show the abundance of mutants with transposon insertion sites within A: the *pur* operon, B: the *cnt* operon or C: the *his* operon. Red is the forward strand and blue is reverse.

Amino acid biosynthesis featured heavily in the dataset. Amino acid utilisation and even auxotrophy has been reported as media dependent (13), and so the substrate availability explains this result. The genes *ilvC* and *ilvD*, involved in branched-chain amino acid biosynthesis (leucine, isoleucine and valine), were both protected during growth in human synovial fluid. The gene which encodes the final step in the biosynthetic pathways for leucine, isoleucine and valine (*ilvE*), however, contained more insertions than expected. Several genes involved in arginine biosynthesis (*argF, argG, argH, arcB*) were also protected, as were *lysC* and *hisA* which are required for lysine and histidine biosynthesis, respectively. Conversely, insertions in genes involved in tyrosine and tryptophan biosynthesis (*aroB* and *aroA*) were beneficial for survival in synovial fluid.

Genes across the *cnt* operon (*cntK, cntM* and *cntE*) were all significantly protected (q<0.05) in synovial fluid (Figure 1) – ie: essential for growth in synovial fluid. These genes encode for the biosynthesis and export of staphylopine, a metallophore believed to play a role in zinc acquisition, identified in *Staphylococcus aureus* (14).

### Using RNASeq to characterise the response of *Staphylococcus epidermidis* to human synovial fluid

In parallel with the TraDIS analysis, to identify genes up- and down-regulated in *S. epidermidis* upon exposure to human synovial fluid, gene expression was studied in two strains (one *ica*-positive and one *ica*-negative) after a 30-minute exposure to either synovial fluid or rich medium using RNASeq. The datasets from both strains demonstrated excellent agreement, with genes identified in both strains (P-adj < 0.05, homology identified using BLAST reciprocal best hits) showing good agreement in levels of changed expression (Figure 2, R^2^ = 0.84). These accounted for 342 upregulated and 263 downregulated genes, with only 11 genes being upregulated in one strain and downregulated in the other. This concordance shows a conserved set of genes respond to synovial fluid exposure. In the dataset generated using *S. epidermidis* RP62A, 341 genes were highly differentially expressed upon exposure to synovial fluid (P-adj. <0.05, log_2_ fold change >1), of which 92 were downregulated and 249 were upregulated. In *S. epidermidis* 846, there were 256 highly differentially expressed genes, of which 64 were downregulated and 192 were upregulated (Supplementary File S1).

**Figure 2:**
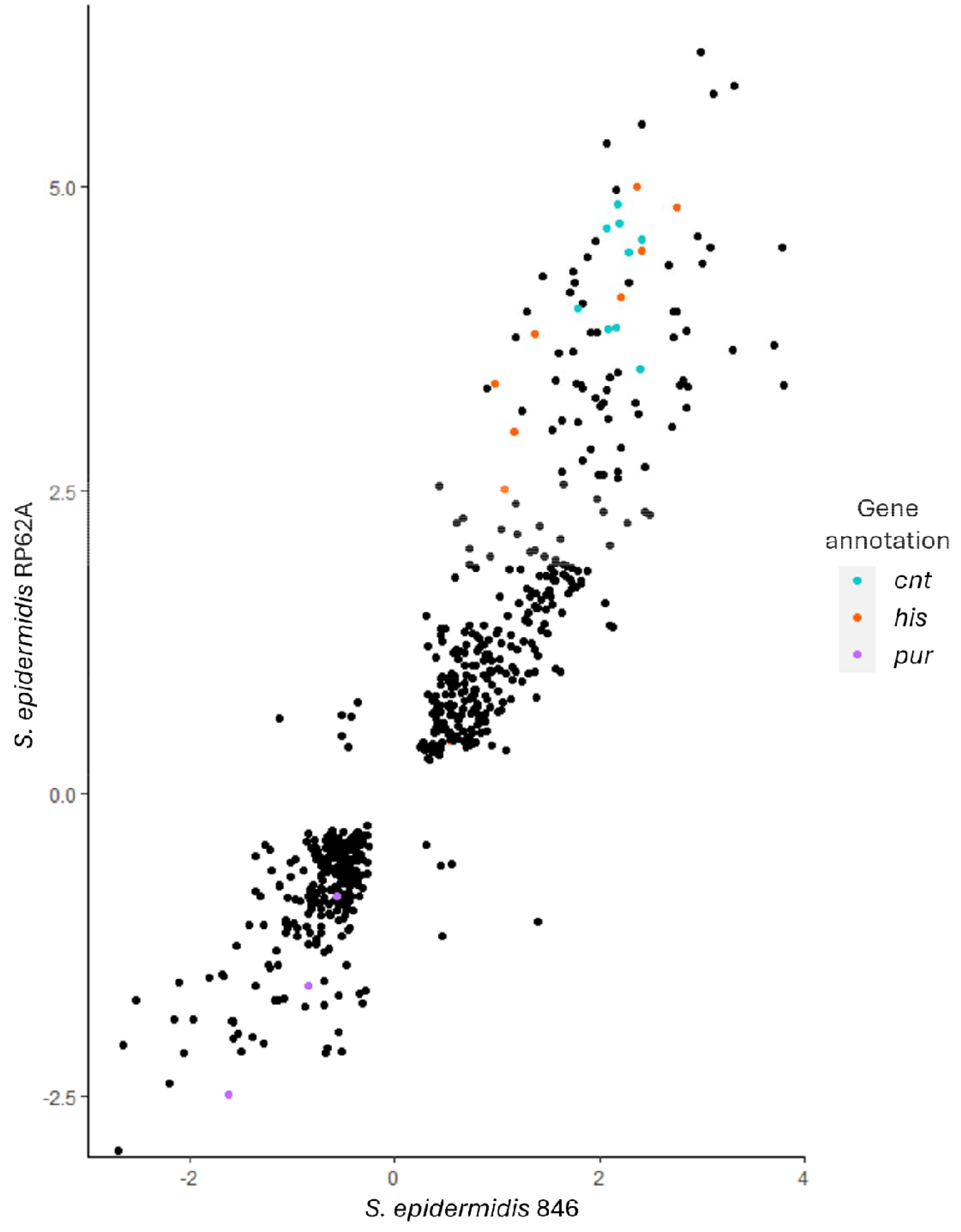
Log_2_ fold change in gene expression after exposure to synovial fluid of genes identified in both strains of S. epidermidis. Genes highlighted belong to key operons. Blue - *cnt*, Orange - *his*, Pink - *pur*.

Generally, genes encoding for amino acid biosynthesis were upregulated, with significant increases in expression of genes for the production of most amino acids. Notable exceptions were cysteine, where the synthase *cysK* was downregulated, and glutamine, where both *rocB* and *pruA* were downregulated. By mapping the genes of interest to the pathways modelled on the biocyc (15) metabolic map for *S. epidermidis* sp. RP62A, significant upregulation of genes encoding for the entire pathway of histidine biosynthesis was observed (Table 2, Figure 2), suggesting that histidine plays a key role in the staphylococcal response to human synovial fluid.

**Table 2:**
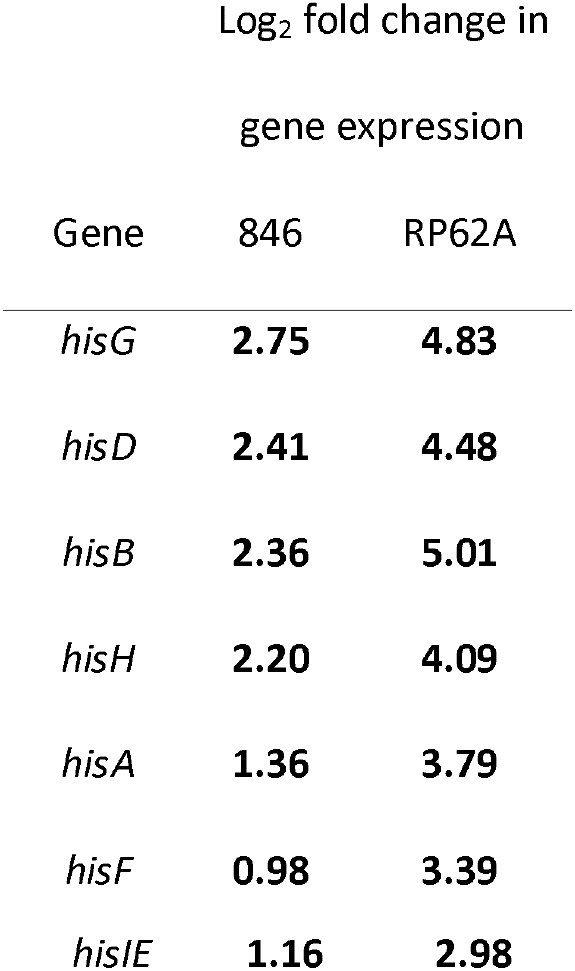
Log_2_ fold change in expression of genes in the *his* genes after exposure of *S. epidermidis* strains RP62A and 846 to human synovial fluid compared to a rich medium control, as quantified using DESeq2. Bold text denotes significant result (p adj < 0.05)

It was noted that differential expression of the *pur* operon was consistent between the two strains, with the first genes in the operon downregulated, genes in the middle not significantly differentially expressed, and the final genes significantly upregulated in RP62A, and demonstrating a positive Log_2_ fold change in 846 (Table 3). When mapping the significant values from RP62A onto the metabolic map, downregulation in the initial part of the inosine-5’-phosphate biosynthesis pathway is observed, with upregulation from the compound 5-amino-1-(5-phospho-D-ribosyl)imidazole-4-carboxamid (AICAR), which is also a by-product of the histidine biosynthetic pathway, as previously mentioned (Figure 3).

**Table 3:**
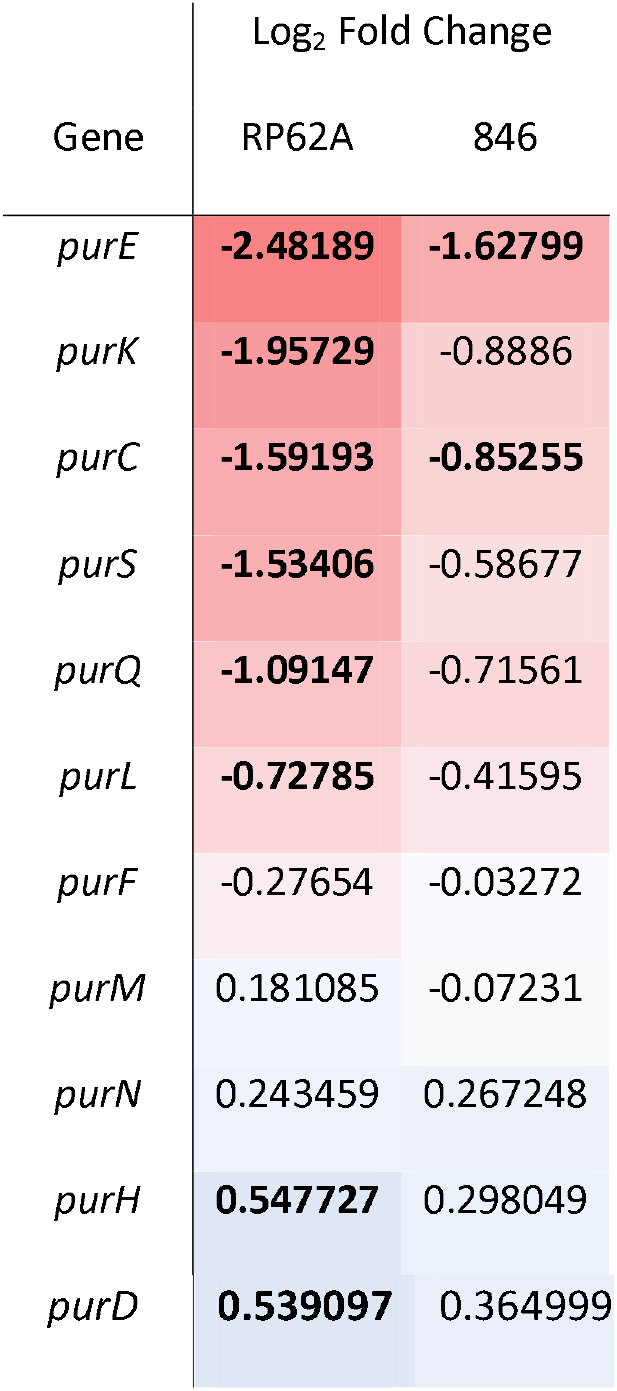
Log_2_ fold change in expression of genes in the *pur* operon after exposure of *S. epidermidis* strains RP62A and 846 to human synovial fluid compared to a rich medium control, as quantified using DESeq2. Bold text denotes significant result (p adj < 0.05)

**Figure 3:**
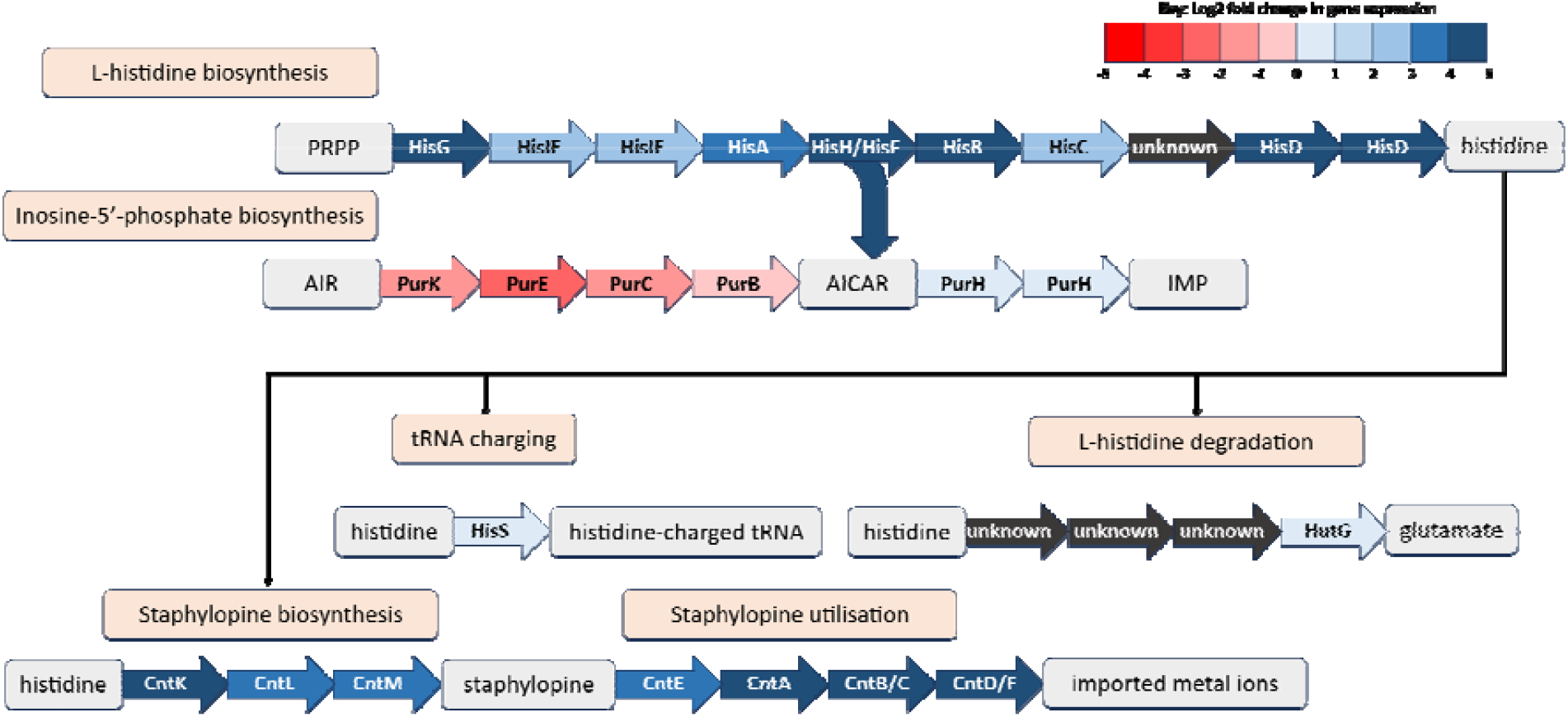
Link between the Pur proteins, His proteins and staphylopine biosynthesis. Blue arrows denote upregulation of the associated gene in the RNASeq data, red arrows denote downregulation, and grey is where the gene encoding this step of the pathway is unknown. PRPP = 5-phospho-α-D-ribose 1-diphosphate. AIR = 5-amino-1-(5-phospho-β-D-ribosyl)imidazole. AICAR = 5-amino-1-(5-phospho-D-ribosyl)imidazole-4-carboxamide. IMP = inosine 5’-monophosphate. Log_2_ fold-changes given are for *S. epidermidis* RP62A after exposure to human synovial fluid.

This link between differential expression of the *pur* operon and the *his* operon warranted further investigation into the how the newly synthesized histidine and inosine 5’-monophosphate (IMP) could be directed. To establish this, the entire dataset of significantly differentially expressed genes (adjusted P <0.05) from *S. epidermidis* was mapped onto the metabolic map hosted on BioCyc (15). This identified multiple upregulated pathways for the utilisation of inosine 5’-monophosphate (IMP), the compound produced by proteins encoded by the upregulated genes in the *pur* operon identified in *Staphylococcus epidermidis* RP62A (Figure 3, Table 2), and three potential pathways for the utilisation of histidine. The pathways for histidine utilisation were i) degradation to glutamate, ii) tRNA charging for protein synthesis, or iii) staphylopine biosynthesis. Of these pathways, staphylopine biosynthesis showed the largest increase in expression, suggesting this may be the driving requirement for histidine (Figure 3).

The proteins required for staphylopine synthesis and utilisation are encoded by the *cnt* operon, where several genes were found to be protected during growth in synovial fluid (relative to growth in a rich medium) in the transposon mutant library experiment (Figure 1). In agreement with this, the entire gene cluster was found to be significantly upregulated in both *S. epidermidis* RP62A and 846 upon exposure to human synovial fluid. Together these data suggest that staphylopine is produced by *S. epidermidis* during growth in synovial fluid.

We then looked at the level of expression of the *cnt* operon within the three RNASeq samples (Figure 4). An expression profile closer to the control was observed in sample 3, which was from a non-infected patient’s synovial fluid sample, whereas the other two used were from cases of prosthetic joint infection. This was further explored using primers designed to amplify the *cntE* gene, which encodes the staphylopine export protein. The *cntE* gene was slightly upregulated in non-infected samples (*S. epidermidis* 846 - not significant, *S. epidermidis* RP62A - *p*<0.05), though the effect was much more pronounced after exposure to synovial fluid from patients with PJI (*p*<0.01 for both strains) (Figure 5).

**Figure 4:**
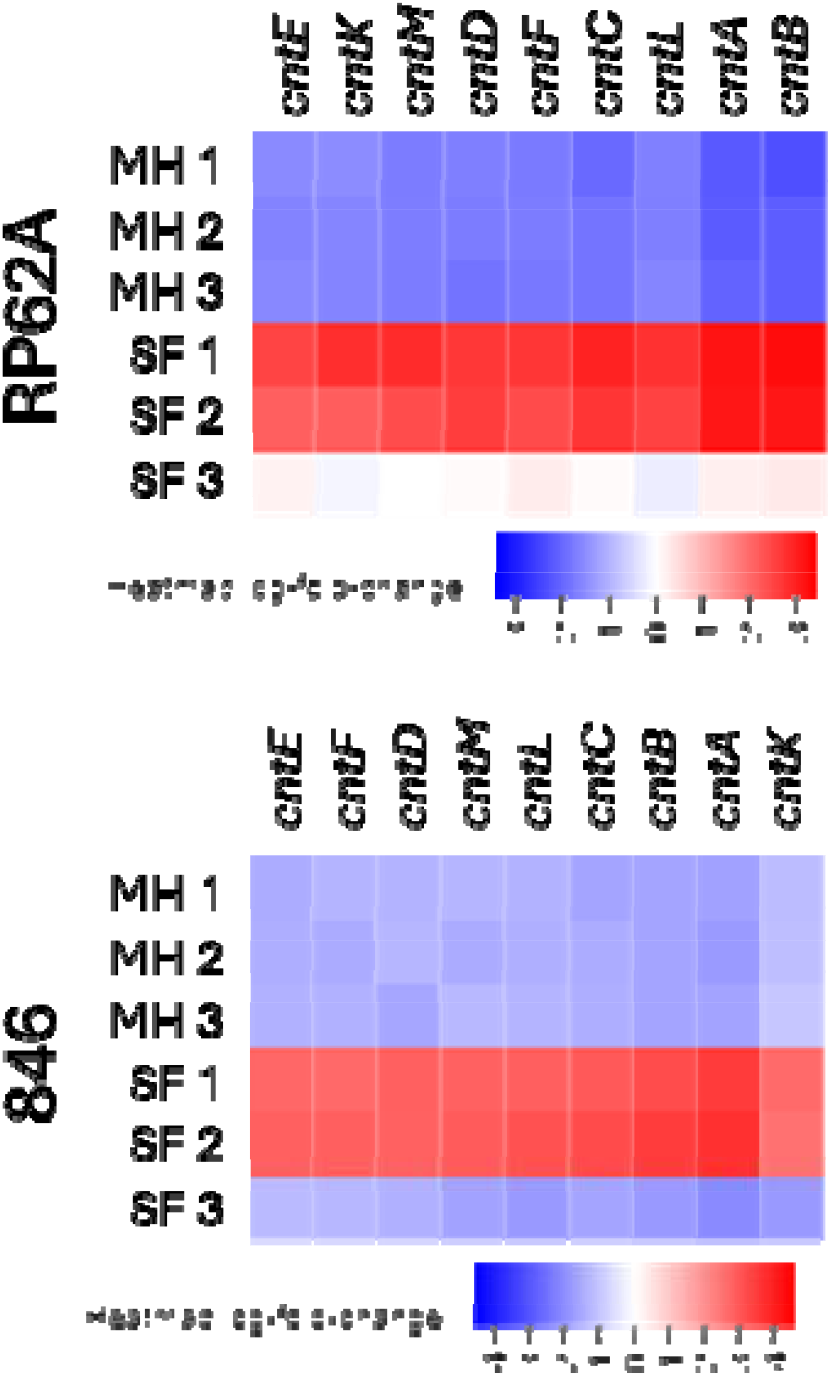
Change in expression of the *cnt* operon after exposure to Mueller Hinton (MH) or Synovial Fluid (SF) in *S. epidermidis* RP62A and 846. Rows are independent biological replicates, with SF1 and SF2 being samples from PJI patients, and SF3 from a patient without PJI.

**Figure 5:**
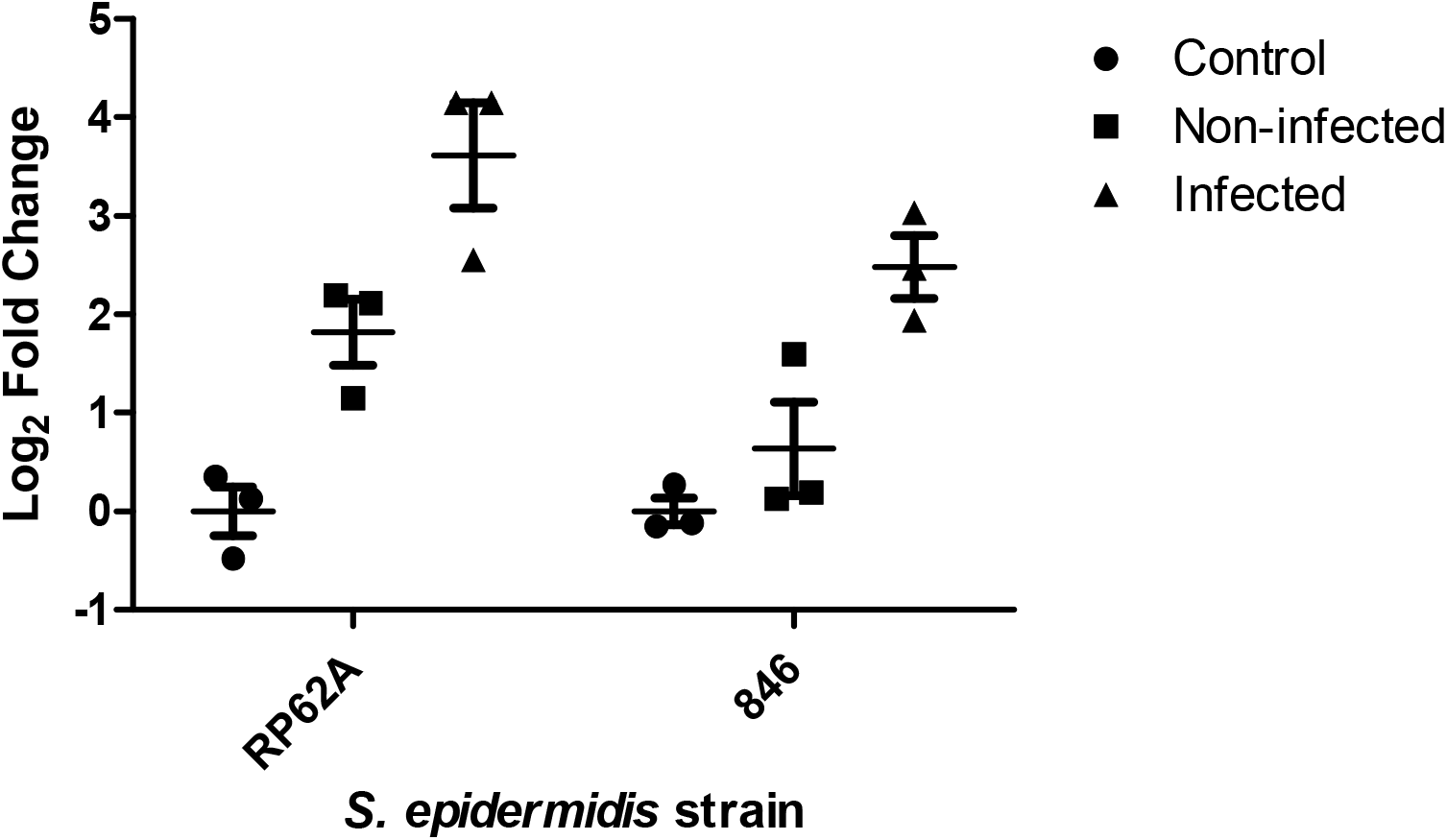
Change in expression of *cntE* in response to processed synovial fluid from either infected or non-infected patients in *S. epidermidis* strains RP62A and 846.

Subsequently, a *cntE* transposon insertion mutant was isolated from the pool using the methods outlined in (16). This isolate demonstrated significantly impaired survival after inoculation into human synovial fluid (not from prosthetic joint infection cases) over 48 hours (Figure 6).

**Figure 6:**
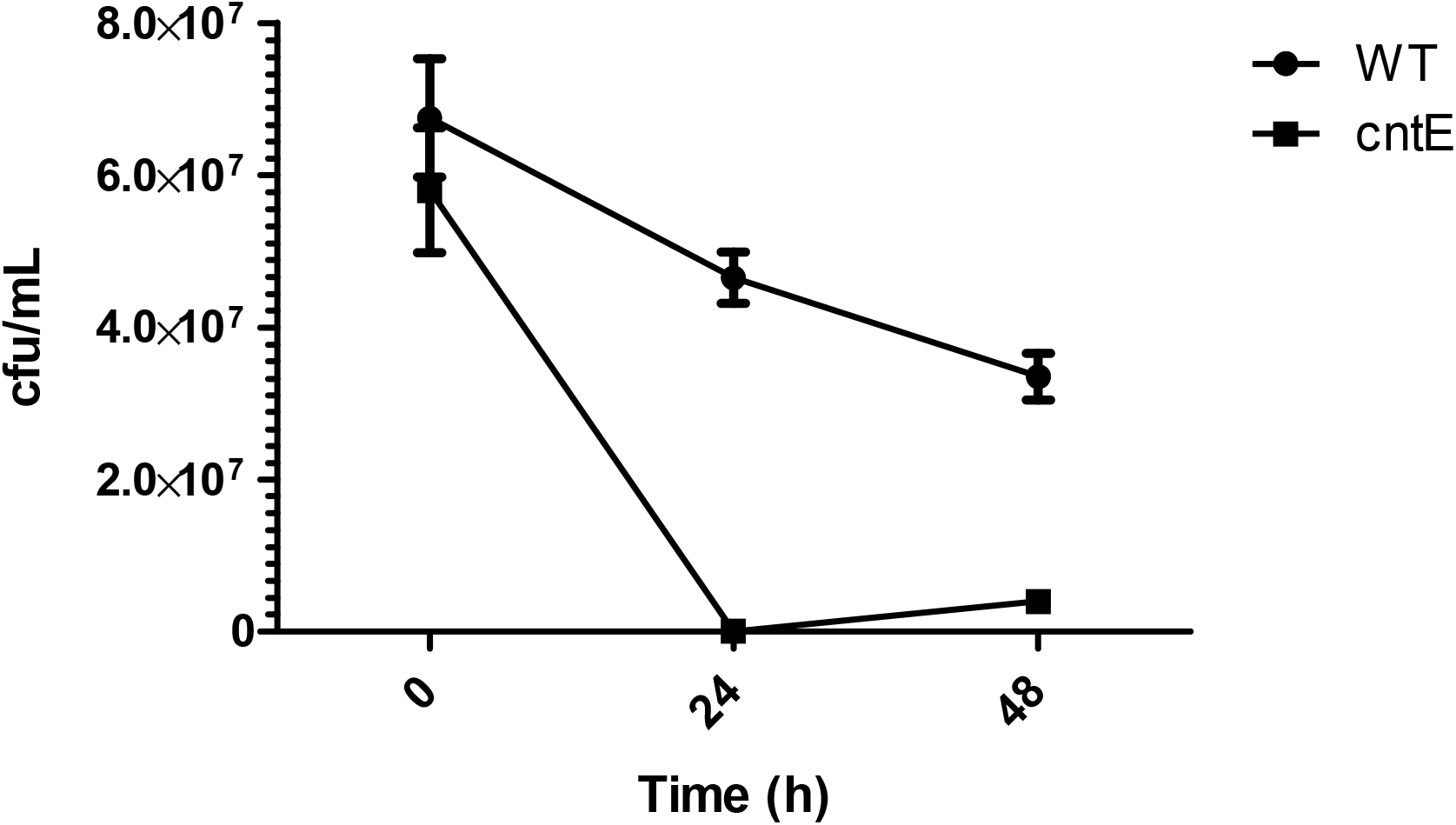
Comparison of survival of *S. epidermidis* 846 wild-type versus a *cntE* transposon insertion mutant in processed synovial fluid over 48h.

Finally, to explore likely conservation of staphylopine production as a mechanism to survive during PJI, the presence of the *cnt* operon was verified in a range of staphylococci isolated from prosthetic joint infection in our previously published collection (17). This included *Staphylococcus epidermidis, lugdunensis, warneri* and *capitis*. Genomic analysis suggested conservation of the *cnt* operon in these staphylococcal species, but it was not found to be present in *Staphylococcus haemolyticus* genomic sequences. Degenerate primers were designed to amplify *cntE*, and tested against a panel of PJI isolates. Two *S. haemolyticus* isolates, which do not encode the *cnt* operon, were included as negative controls, and a model *S. aureus* NCTC 8325 was included due to the high proportion of PJIs caused by the species (Figure 7). The primers successfully amplified *cntE* from DNA extracted from all of the non-*haemolyticus* PJI isolates, suggesting that these may provide a useful tool to aid PJI diagnosis by identifying the presence of the *cntE* gene from bacterial nucleic acids extracted from synovial fluid samples, though further work to optimise this would be required.

**Figure 7:**
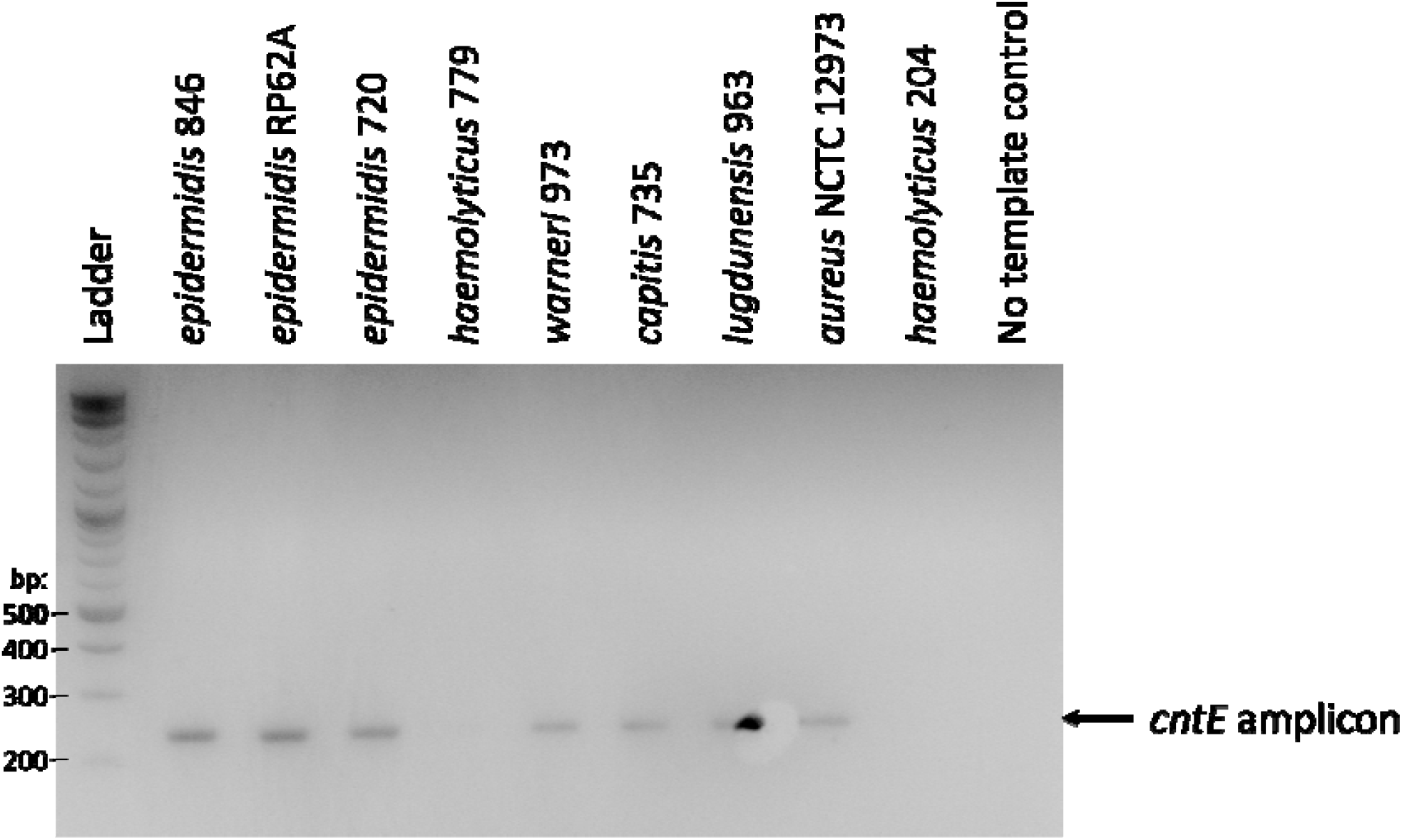
Amplification of *cntE* from a panel of staphylococci using degenerate primers. *S. haemolyticus* does not contain the *cnt* operon.

## Discussion

Diagnosis of prosthetic joint infection remains difficult when the causative agents are coagulase-negative staphylococci, and their ability to form biofilms reduce treatment efficacy (8-10). To better understand the mechanisms utilised by *Staphylococcus epidermidis* to survive in human synovial fluid, and how they respond to it, we used a combined approach of a genomic fitness screen with a transposon mutant library and studied the transcriptome using RNASeq. Whilst several methods of synthetic synovial fluid production are available (18-20), and can be used to replicate the aggregating behaviour of staphylococci in human samples (19), we used human samples from patients diagnosed as possible PJI to most closely mimic *in vivo* conditions, as previous research has demonstrated a difference in growth and biofilm composition based on the medium available (21).

Two *Staphylococcus epidermidis* strains were selected for RNASeq analysis (one well characterised *ica*-positive (RP62A) and one *ica*-negative (846)). These offered the opportunity to investigate which biofilm production methods may be utilised during PJI. We noted no upregulation of *icaA* after exposure to human synovial fluid, which is vital for polysaccharide intercellular adhesin (PIA) based biofilm formation, which has been proposed as a target to identify biofilm-forming strains of staphylococci, though discounted as a useful diagnostic marker for PJI (22). Conversely, proteinaceous biofilms utilise a range of proteins, of which genes encoding both the accumulation-associated protein (*aap*) and the *S. epidermidis* autolysin *atlE* were upregulated in both strains tested (*aap*=SERP2398, +1.14, BHIJLP_01729, +0.88. *atlE*=SERP0636, +0.99, BHIJLP_00991, +0.62(not significant)), though neither were identified as playing a role in survival in the transposon mutant library. Our data suggest the preference for proteinaceous biofilm formation upon exposure to synovial fluid, which may mediate the aggregate formation observed upon exposure to synovial fluid of several staphylococcal species (6, 18, 19).

The *pur* and *his* operons identified in both the transposon insertion sequencing and RNASeq experiments have also been linked to biofilm formation. The purine biosynthesis pathway encoded by the *pur* operon is important for the production of eDNA, a major component of the biofilm matrix of *S. aureus* (23), which was upregulated during biofilm formation of *S. aureus* strains (16). Conversely, we identified downregulation of the genes situated early in the operon, with downstream increased expression of *purH* only being significant in *S. epidermidis* RP62A. This is possibly due to the increased levels of histidine biosynthesis, which lead to production of the compound 5-amino-1-(5-phospho-D-ribosyl)imidazole-4-carboxamide (AICAR) which feeds into purine biosynthesis using *purH* (Figure 3) and possibly has a feedback effect on the genes upstream of AICAR. This interplay between the metabolic pathways has been previously demonstrated, where *S. aureus pur* mutants could still produce IMP via the histidine biosynthetic pathway (24).

We recently identified *hisB* as a marker of staphylococcal biofilm formation (12), and a *Staphylococcus xylosus hisB* mutant has previously been demonstrated reduced biofilm formation (25). This protein enables a step in histidine biosynthesis after the diversion of AICAR to the purine biosynthetic pathway, suggesting the link is between histidine biosynthesis and biofilm production, rather than an indirect link to purine biosynthesis via AICAR. Prior research has identified reduced histidine levels in infected synovial fluid compared to non-infected (26), suggesting infectious organisms may both produce their own and import histidine from the host. Histidine import has recently been shown to play an important role in acid tolerance in *Staphylococcus aureus* (27) using the histidine permease designated 0846. Homologues of this were found and significantly (p-adj <0.05) upregulated in both of our strains after exposure to synovial fluid (BLJHFILP_01093, +0.68; SERP0529, +1.33), suggesting that they are both importing histidine. This gene was not protected in the TraDIS dataset during growth in synovial fluid, though this may be due to a preference for biosynthesis over import.

Histidine can be utilised by bacteria in a variety of ways. It can be used to produce glutamate via HutG, which can act as an important nitrogen source (28), or to charge tRNAs for protein production using HisS. The genes encoding both of these proteins were found to be significantly upregulated in *S. epidermidis* RP62A after exposure to synovial fluid (Log_2_ fold change +0.42 and +0.43 respectively), whereas only *hisS* was upregulated in *S. epidermidis* 846 (+0.53). This of interest as glutamate is present in synovial fluid and so bacterial production of glutamate should not be necessary (29), suggesting a possible overspill linked to the increase in histidine production. Histidine is also utilised by CntK for the production of the metallophore staphylopine (14). The entire *cnt* gene cluster (*cntKLMABCDFE*) encodes for the production and utilisation of the metallophore (14), and we found it to be entirely upregulated to a high level in both strains (Figure 2). In combination, genes from the cluster were protected during growth of *S. epidermidis* 846 in synovial fluid (Figure 1), suggesting that staphylopine plays a key role in both the immediate response to and longer-term growth in synovial fluid, which was validated with a defined mutant. These combined data suggest this is likely the pathway which the excess of imported and produced histidine is required for.

Staphylopine has been studied extensively in *S. aureus* as an important compound capable of outcompeting the human protein calprotectin in zinc acquisition (30). Calprotectin is a biomarker used in the diagnosis of PJI (31); however the use of host markers as a diagnostic tool is more complex for infections caused by CoNS, which often yield false-negative results (32), making a positive bacterial biomarker an attractive opportunity for novel diagnostics.

The *cnt* genes have been previously identified in *S. epidermidis, cntL* and *cntA* were found to be involved in multistress responses, proposedly due to an increased requirement for metal-containing proteins under stress conditions (33). Metal acquisition has been identified as playing a key role in the *Staphylococcus* response to human fluids, though this is mostly focused on *S*.*aureus*. Upregulation of genes encoding iron transport proteins has been observed in *S. aureus* after exposure to human blood (34), and in *S. aureus* removed from PJI samples, where *cntAKLM* were also upregulated, though not to the same high level as many other genes linked to iron acquisition (35). The transcriptome of *S. aureus* has also been studied in infected mouse livers, where a wide range of metal transporters were upregulated 24 h post-infection (36). Whilst *cnt* was recognised, and a *cntE* and *cntK* mutant demonstrated reduced virulence, a much larger difference in expression was observed for *mntABC*, which was not identified in our work. Iron uptake genes have also been linked to the *S. epidermidis* biofilm transcriptome after exposure to human blood, though the *cnt* system was not identified (37). More recently, RNA has been extracted from sonicates of prosthetic joints removed due to *Staphylococcus epidermidis* infection. In these samples, the *cnt* genes were also expressed to high levels (38), suggesting that our findings are reflected *in vivo*.

Here, we have clearly demonstrated that an insertional mutation in the staphylopine export protein, *cntE*, compromises *S. epidermidis* survival in synovial fluid, further validating the theory that staphylopine is vital for staphylococcal survival *in vivo* during PJI. The high level of conservation we observed across multiple staphylococcal species suggests this is a key mechanism of metal acquisition across the genus. *S. haemolyticus* is an exception, where investigation of other metal-acquisition proteins may be of future interest, though it should be noted this species accounts for a relatively small proportion of PJI cases (39). The presence of the *cnt* genes across the majority of staphylococci causing PJI opens the opportunity for this to be exploited as a novel diagnostic tool.

## Methods

### Acquisition and processing of synovial fluid samples

Synovial fluid samples were collected by the Norwich Biorepository from the Norfolk and Norwich University Hospital as diagnostic or therapeutic excess under ethical approval (Reference ID ETH2122-1102; Norwich Biorepository licence NRES number 19/EE/0089; IRAS Project ID 259062). Samples were anonymised, and processed to render them acellular: The samples were centrifuged to remove cells and debris, followed by dilution 1:1 with PBS containing 0.1% glucose. This enabled filter sterilisation using 0.22 um pore size syringe filters whilst maintaining the glucose concentration to provide a carbon source. PBS was selected as it would aid in maintenance of the pH during growth, which would be achieved *in vivo* with the constant turnover of synovial fluid (40). The addition of 0.1% glucose as a carbon source was based on this being within levels observed in human synovial fluid samples (41). Processed samples were stored at −80 °C until required.

### Production of a transposon mutant library in *Staphylococcus epidermidis* 846

The pIMAY plasmid (Addgene) (42) was selected due to the ability to counter-select using a *Staphylococcus* specific heat treatment or induction of a kill gene (anti-*secY*). The plasmid was modified by adding a Tn5 transposon containing an erythromycin resistance cassette and a gene encoding Tn5 transposase, synthesized by IDT technologies (Supplementary File S2). The plasmid was maintained in *S. aureus* DSM-23609 at 28 °C, then extracted using the Qiagen plasmid mini prep kit according to manufacturer’s instructions. A panel of coagulase-negative *Staphylococcus* isolates from our previous work (12, 17) were tested for their electrocompetency using this plasmid. *Staphylococcus epidermidis* 15TB0846 was found to be the most competent (efficiency 10^1^ transformants/μg DNA), and therefore carried forward for the production of a transposon mutant library.

Competent *S. epidermidis* 846 cells were produced by diluting an overnight culture 100-fold in fresh BHI, then incubating at 37 °C with shaking to an OD_600_ of 0.5-0.9. A total volume of 3 mL culture per electroporation was used, which was harvested by centrifugation (4 °C, 3000 ×*g* 10 min), washed 3 times in ice cold sterile water, twice in ice cold sterile glycerol and finally resuspended in 100 μL ice cold sterile 10% glycerol. Purified pIMAY:Tn5 was used to transform the cells using electroporation (0.1 cm gap cuvette, 1800 V, 25 μF, 100 Ω). Initial transformants were grown in BHI at room temperature over 72 h to increase the library diversity and integration frequency, then subjected to plasmid removal in a two-step protocol. Firstly, the cells were heated to 42 °C for 72h, then grown in BHI containing 5 μg/mL erythromycin to remove any remaining parent cells. The potential transformants were then grown in BHI containing 15 μg/mL erythromycin and 1.25 μg/mL anhydrotetracycline (Sigma) to remove residual plasmid. Approximately 95% of plasmid was confirmed to have been cleared from the transposon mutant library by comparing growth of serial dilutions on BHI containing erythromycin (5 μg/mL) or erythromycin (5 μg/mL) and chloramphenicol (10 μg/mL).

Growth of the transposon mutant library to investigate genes essential for growth in synovial fluid: An aliquot of library containing 3 × 10 ^7^CFU was inoculated into 2 mL of either processed synovial fluid or Müller Hinton (MH) broth and grown at 37 °C for 48 h. Cells were harvested by centrifugation and washed twice with PBS, then the centrifugation repeated. The pellets were resuspended in lysis buffer (20 mM Tris-HCl, 2 mM EDTA, 1% SDS, 1.2% Triton X-100, pH 8.0) containing 0.5 mg/mL lysostaphin, and incubated at 37 °C for 30 min to lyse the cells. DNA was then extracted using the Zymo Quick-DNA miniprep kit according to manufacturer’s instructions.

### Library preparation, sequencing and data analysis

Genomic DNA extracted from the transposon mutant library was diluted to 11.1 ng/µL and tagmented using MuSeek DNA fragment library preparation kit (ThermoFisher, USA). Fragmented DNA was purified using AMPure XP (Beckman Coulter, USA). DNA was amplified by PCR using biotinylated primers specific to the transposon and primers for the tagmented ends of DNA. PCR products were purified again using AMPure XP beads and incubated for 4 h with streptavidin beads (Dynabeads) to allow for capture of the DNA fragments with the transposon. A subsequent indexing PCR step using barcoded sequencing primers allowed for the pooling of samples. Streptavidin beads were magnetically removed from the PCR products which were further purified and size-selected using AMPure XP beads. The indexed library was quantified using Qubit 3.0 (Invitrogen, USA) and Tapestation (Agilent Technologies, USA). The library was sequenced using NextSeq 500 Illumina machine with a NextSeq 500/550 High Output Kit v2.5 (75 cycles) (Illumina).

Results were analysed using BioTraDIS (version 1.4.1) (43). BioTraDIS matched sequence reads against the reference genome (Supplementary File S3) using BW aligner and created insertion plots of mapped transposon insertion sites. These results identified the presence of approximately 57,000 unique insertion mutants within the library.

To determine essentiality, the R package within the BioTradis software was used, version 1.4.5 (43). The insert plot files generated from the pipeline were combined with the annotated reference and the tradis_gene_insert_sites command aligned the insertion sites to genes. Then, the tradis_essentiality.R command determined gene essentiality and output three .csv files per sample: one for essential genes, one for ambiguous genes, one for nonessential genes.

For comparison, the relevant gene insertion site files were grouped and added to either a control or condition list. These lists were provided to the script tradis_comparison.R that uses EdgeR to identify genes with differences in insertion density.

Genes were considered to be involved in survival of the strain in synovial fluid if the adjusted p value (q) was below 0.05, and the Log_2_ fold change reported was above 1 or below −1.

### RNA extraction for RNASeq and RT-qPCR

Overnight cultures of *Staphylococcus epidermidis* strains RP62A and 846 were diluted 1/100 into 10 mL MH broth and incubated with shaking until an OD_600_ of 0.2-0.3 was reached. Cells were harvested by centrifugation and washed in PBS before resuspending in 500 μL of either processed human synovial fluid or MH broth for 30 min at 37 °C. Cells were then recovered by centrifugation, washed twice in PBS and resuspended in 800 μL Zymo DNA/RNAShield. The cell suspension was processed using the lysing matrix from Zymo fungal/bacterial Quick-RNA miniprep kit in an MP Fastprep at 6.5 m/s; 1 min on, 5 min off repeated five times. The samples were then processed using the kit according to manufacturer’s instructions, with the addition of the optional on-column DNase treatment. DNA and RNA concentrations of the resulting samples were quantified by Qubit (Thermo Fisher), sample quality was assessed using a combination of Nanodrop and TapeStation (Agilent) electrophoresis, where RIN values were confirmed to be above 6. Library preparation and sequencing were performed by Azenta (Germany).

### RNASeq data analysis

RNAseq data was analysed using the same tools outlined in (44) which were run using Galaxy (45). Briefly, read quality was assessed using fastQC (46), followed by sequence trimming with fastp (47). Reads were aligned to the reference genomes (*S. epidermidis* RP62A GenBank assembly CP000029.1; *S. epidermidis* 846 Supplementary File S3) using HISAT2 (48), and the number of reads successfully mapped quantified using samtools stats (49). Mapped reads were matched to annotated regions using featurecounts (50), and differential gene expression quantified using DESeq2 (51). Genes were considered differentially expressed if the adjusted P-value (p-adj) was below 0.05 and the Log_2_ fold change was above 1 or below −1.

### RT-qPCR

RT-qPCR primers were designed using Primer3 software (v4.1.0) (52) (Table 4). Primers were validated using a standard curve of DNA concentrations varying from 0.06 to 2 ng per 10 μL reaction. The presence of single products was checked using visualisation of the dissociation curves. RT-qPCR was performed in 10 μL reactions using NEB Luna One-Step RT-qPCR according to manufacturer’s instructions. Gene expression in synovial fluid was calculated versus the media controls, using *rpoB* as a housekeeping gene and the 2 ^ΔΔCT^ method.

**Table 4:**
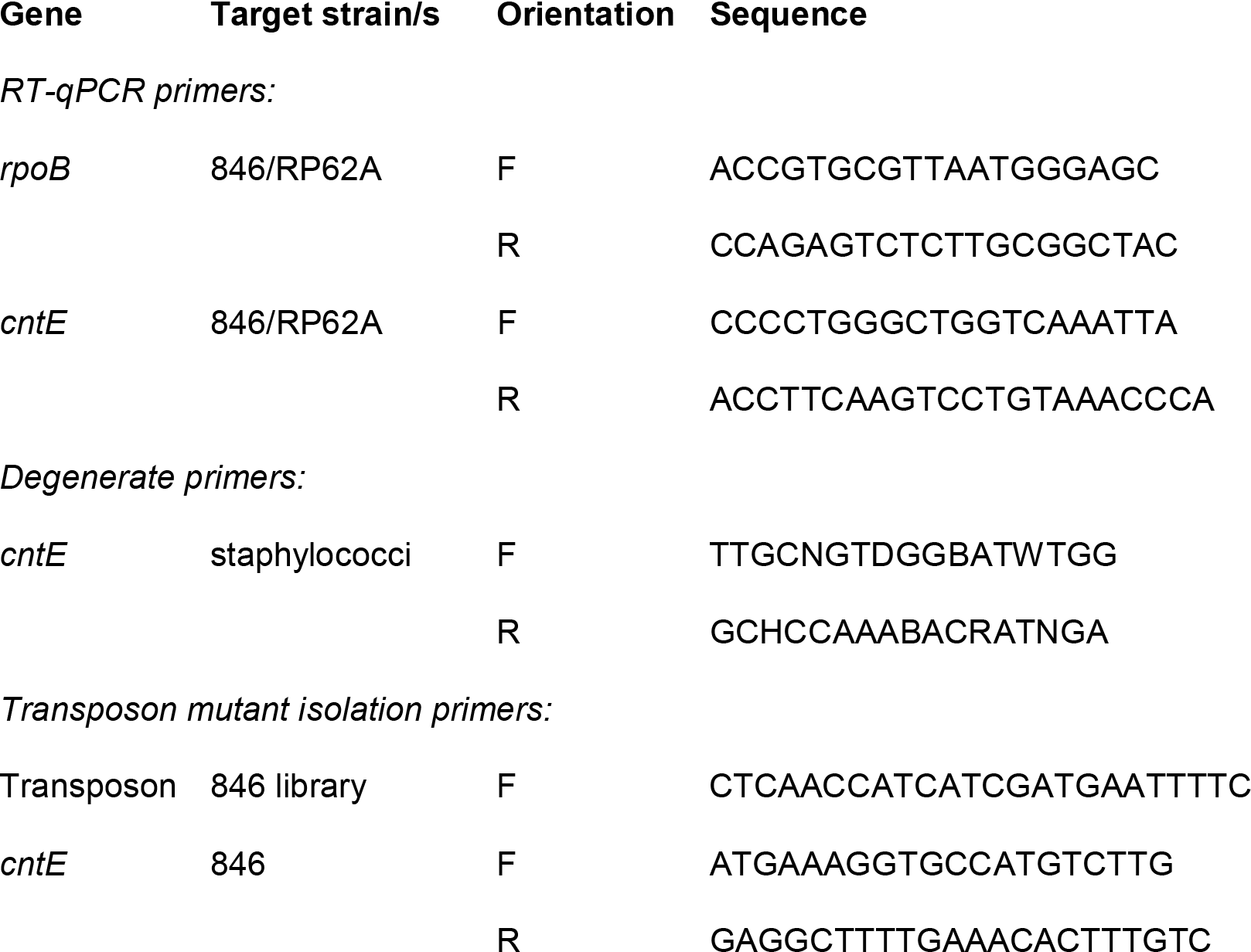
Primers used in this study.

### Isolation of Tn5::*cntE* mutants from the TraDIS pool

Specific transposon insertion mutants were isolated from the TraDIS pool using PCR. A forward primer was designed to bind the transposon, after the mosaic end, and reverse primers to the end of the genes of interest (Table 4). The transposon mutant library was diluted 1/10^6^ and 200 μL aliquots were grown at 37 °C for 48 h in 96-well plates. Rows were pooled and concentrated 10x before the concentrated culture was added to a 40-cycle PCR reaction using Promega G2 GoTaq according to manufacturer’s instructions (2.5 μL culture/25 μL reaction). Another PCR was performed on individual wells from the row of interest to identify which contained the potential mutant. This process was repeated on the target well with a 10x higher dilution each time, until the initial dilution factor was 10^9^. At this point the well of interest was diluted by 10 ^6^before plating onto Mueller-Hinton agar and growth overnight at 37 °C. Colony PCR was performed using the same parameters, and finally colonies of interest were subjected to colony PCR with primers which amplified the gene of interest only, and compared to DNA extracted from the wild-type to confirm the presence of inserted transposon.

### Quantification of mutant growth in processed synovial fluid

Mutants were assessed for their ability to grow in processed synovial fluid. Overnight cultures were inoculated into MH broth and incubated at 37 °C for 16h. The resulting cells were washed twice in an equal volume of PBS before being used to inoculate processed synovial fluid samples using a 1/100 dilution in 200 μL cultures in a 96-well plate. Samples of initial inoculum were removed, and cultures were harvested for cfu quantification at 24 h and 48 h. For each, serial dilutions were performed in PBS and cfu counts were performed on MH agar.

## Supporting information

Supplemental analysis output files

## Supplementary Information

### Excel file

S1 File. Significant (p-adj <0.05) results from RNASeq analysis (DESeq2 output) and TraDIS analysis (BioTraDIS output)

- 846 DeSeq2 output (significant only)
- 62A DeSeq2 output (significant only)
- BioTraDIS output (significant only)

S2 File. Tn5 transposon synthesized by IDT technologies.

## Data access statement

The short read sequences generated in this project are available from the NCBI Sequence Read Archive (SRA) under BioProject PRJNA1272027.

## Acknowledgements

We would like to acknowledge the Norwich Research Park Biorepository for their role in sample collection, anonymization, and storage.

EM, CH, HF, JW and IM were supported by a grant awarded to JW and IM by Heraeus Medical. LS was supported by funding awarded to MW and IM by Action Arthritis, the Norfolk and Norwich University Hospital charity, and Quadram Institute Bioscience. MY and MW were supported by the Biotechnology and Biological Sciences Research Council (BBSRC) Institute Strategic Programme Microbes and Food Safety BB/X011011/1 and its constituent project BBS/E/F/000PR13636. The funders had no role in study design, data collection and analysis, decision to publish, or preparation of the manuscript.

